# Development of a novel loop-mediated isothermal amplification (LAMP) assay for *Balamuthia mandrillaris*

**DOI:** 10.64898/2026.09.18.752794

**Authors:** Chenyang Lu, Chayan Sharma, Shreyes Kanumuru, Christopher A. Rice

## Abstract

*Balamuthia mandrillaris* is a free-living amoeba that can cause cutaneous skin lesions, systemic disease, and a severe life-threatening CNS infection, *Balamuthia* amoebic encephalitis (BAE), which has a high mortality rate (~92%). The current diagnostic methods for *B. mandrillaris* have many limitations including the long detection time, low sensitivity, and specific diagnostic tests only available in highly specialized centers. The early stage of diagnosis is important for the treatment because of how quickly BAE infection then develops. Therefore, we developed, optimized, and evaluated a novel loop-mediated isothermal amplification (LAMP) assay for the 18S rRNA gene of *B. mandrillaris*, which can be observed by naked eyes with phenol red. This novel diagnostic assay can detect ten different *B. mandrillaris* isolates with no cross-reactivity with the DNA of other free-living amoeba, protozoa, fungi, or bacteria tested. Compared with conventional PCR, our LAMP assay displayed ~10-100-fold higher sensitivity from extracted DNA, heat-treated samples, and non-treated samples, with the limit of detection (LOD) of 10 fg/μL DNA, 10 trophozoite/100 μL for non-treated cells, and 1 trophozoite/100 μL for heat-treated cells, respectively. This method was further validated using two newly developed qPCR and dPCR methods to determine the LOD of approximately 1 copy/μL. In addition, there is no tolerance for complicated matrices, that LAMP maintained efficacy in cerebrospinal fluid (CSF), urine, and blood-spiked samples. Due to the simplicity, robustness, and high sensitivity of the developed LAMP method, it will be useful for quick clinical detection and diagnosis of *B. mandrillaris*, particularly in non-specialized centers and resource-limited areas. Here we report the first LAMP diagnostic method for *B. mandrillaris*; publicly disclosed on Tuesday, November 4^th^, 2025 at the 20th (XX) International Meeting on the Biology and Pathogenicity of Free-Living Amoebae (FLAM 2025) at Puerto Morelos, Mexico.

## Introduction

*B. mandrillaris* is a free-living amoeba that causes *Balamuthia* amoebic encephalitis (BAE), a “rare” but fatal central nervous system infection in humans and animals (Visvesvara 2007). The clinical diagnosis of BAE remains extremely challenging, as early symptoms are often subtle or non-specific, and disease suspicion frequently considers tumors, abscesses, other inflammatory neurological conditions or more common meningitis causing pathogens. In the early clinical stage, the conventional diagnostic methods contain neuroimaging methods (like CT and MRI) and histopathological examination. The lack of specificity for BAE often lead to misdiagnosis and many cases are diagnosed only postmortem (Khurana 2022). Immunohistochemical staining and immunofluorescence assays are necessary for the diagnosis, but these tests need specific antibodies and are only available in a few laboratories and research centers (Cope 2019). In some geographical areas, like China and Peru, many patients with BAE initially present with cutaneous lesions prior to central nervous system involvement exhibit significantly higher survival rates due to earlier clinical recognition and intervention (Chen 2025), demonstrating the importance of earlier diagnosis in these patients. In the USA, we typically see patients present with BAE disease only without cutaneous lesions. Speculating that different geographical genotypes or strains have differing pathobiology mechanisms similar to different *Leishmania* species.

Molecular diagnostic methods are more specific, some PCR-based assays have been developed (Wang 2020, Yagi 2005, Kiderlen 2008, and Ahmed 2011). These molecular approaches have demonstrated efficacy in clinical and environmental samples with high specificity and sensitivity. However, these tests are hypothesis-driven and typically used in cases where *B. mandrillaris* infection is already suspected based on clinical symptoms. Next-generation sequencing (NGS) technology is an excellent identification method of unknown pathogens. This high-throughput, unbiased, diagnostic method can detect thousands of microorganisms, making it particularly useful for identifying rare or unexpected pathogens such as *B. mandrillaris* or other FLA (Feng 2025 and Sakiyama 2025). NGS techniques are always influenced by low pathogen load and host nucleic acid background, especially in *B. mandrillaris* cases, where CSF is commonly tested but often contains low-to-no pathogen load, so repeated sampling and sequencing are necessary for confirmation. Although NGS-based diagnostics are used if the initial diagnosis fails, typically this is delayed by a few days to receive the initial results; and their high cost and technical complexity to run the instrumentation restricts the widespread clinical application as a first-line diagnostic method.

Loop-mediated isothermal amplification (LAMP) was first described in 2000, it is a nucleic acid amplification technique that is rapid, sensitive, and highly specific (Notomi). The LAMP assay typically contains a set of six primers, including two outer primers, two inner primers, and two loop primers. Amplification proceeds under isothermal conditions, so it can be performed using simple equipment such as a water bath or heating block, and amplification results can be directly visualized by colorimetric changes with the naked eye, without the need for electrophoresis or advanced visualization instrumentation. Compared with conventional PCR, LAMP exhibits higher analytical sensitivity, often exceeding PCR by 10- to 100-fold (Park 2022). The reduced equipment requirements and simplified workflow lower operational costs and make LAMP suitable for deployment in primary healthcare facilities, remote regions, and field-based epidemiological investigations (Picot 2020). LAMP has already been applied for the detection of other free-living amoeba; for example, Mahittikorn et al., developed a LAMP assay targeting a virulence-related gene of *N. fowleri*; and two LAMP assays targeting 18S rRNA gene of *Acanthamoeba* (Yang 2013, Aykur 2023, and Ge 2013). However, there is no previously developed or published LAMP assay specific for *B. mandrillaris*.

Therefore, the development of a robust and reliable LAMP diagnostic assay for *B. mandrillaris* can improve early detection and potentially reduce mortality rates. Furthermore, this assay could be used for environmental sampling and monitoring to understand geographical distribution, identify populations or areas that are at higher risk of exposure, and support future public policies to protect individuals from becoming patients. In this study, we aimed to develop and validate a rapid, specific, and sensitive LAMP diagnostic assay targeting 18S rRNA gene for *B. mandrillaris*, and to systematically evaluate its specificity, sensitivity, and performance in both laboratory and clinically relevant samples.

## Materials and Methods

### Amoeba Culturing

All *B. mandrillaris* isolates [V039, ITSON (donated by Dr. Luis Fernando Lares-Jiménez, ITSON University, Mexico), SAM, RP5, OK1 (donated by Dr. Thelma H. Dunnebacke, California Department of Public Health, US), V433, V188, V194, V426, and V619 (ordered from BEI resources, NIH supported program, US)]. Other free-living amoeba used in the study were American Type Culture Collection (ATCC) strains *Naegleria fowleri* strain Nf69 (ATCC 30215) and *Acanthamoeba castellanii* genotype T4 (ATCC 50370). All *B. mandrillaris* were maintained at 37°C, 5% CO_2_ in *Balamuthia mandrillaris* ITSON medium (BMI) (Lares-Jiménez 2015), supplemented with 10% fetal bovine serum (FBS; Corning, NY, USA) and 125 µg 5000 U/mL penicillin /5 mg/mL streptomycin (pen-strep; Gibco, MD, USA) in vented 75 cm^2^ tissue culture flasks (Genesee Scientific, CA, USA). *N. fowleri* was grown axenically at 34°C in Nelson’s complete medium (NCM), supplemented with 10% FBS and 125 µg pen-strep, in non-vented 75 cm^2^ tissue culture flasks (Genesee Scientific, CA, USA) (Rice 2015); *A. castellanii* was grown axenically at 27°C in Protease Peptone-Glucose Media (PG), supplemented with 125 µg pen-strep in non-vented 75 cm^2^ tissue culture flasks. Routine subculturing was performed every 3 to 4 days when the cells were between 80-90% confluence. For passaging cells, 5 mL 0.25% Trypsin-EDTA (1X) (Gibco, MD, USA) was added to the flasks containing *B. mandrillaris* and incubated for 5 min at 37°C to detach these cells. *N. fowleri* flasks were placed on ice for 15 min and *A. castellanii* cells were mechanically harvested to detach the cells from the flasks. The cells were collected in a 50 mL centrifuge tube and then centrifuged at 3,214 x g (3,900 rpm) at 4°C. Cells were enumerated, diluted to the appropriate concentration and sub-cultured for continuing their growth.

### Sample Preparation

*Sarcocystis neurona* was provided by Dr. Sriveny Dangoudoubiyam (Purdue University); *Toxoplasma gondii* was provided by Dr. Christoph Konradt (Purdue University); *Candida auris* and *C. albicans* were provided by Dr. Shankar Thangamani (Purdue University). *Escherichia coli* (ATCC 8739), *Serratia marcescens* (ATCC 13048), and *Pseudomonas aeruginosa* (ATCC 9027) were bought from ATCC.

Genomic DNA was extracted from all organisms using a commercial DNA extraction kit (QIAGEN, MD, USA), according to the manufacturer’s instructions. Each DNA sample was extracted from 2×10^6^ cells. For *B. mandrillaris* isolate V039, purified genomic DNA was serially diluted in molecular grade nuclease-free water (dH_2_O; Corning, NY, US) to generate a range of DNA concentrations from 0.1 fg/μL to 100 pg/μL. Heat-treated and original (untreated) *B. mandrillaris* trophozoite samples were also prepared for the downstream analysis. Briefly, the cell pellet was counted and diluted in nuclease-free water to a final cell number of 10,000 trophozoites/mL, and then a 10-fold serial dilution was performed six times. *B. mandrillaris* were heated at 95 °C for 1 h and the original cells were directly used without further processing for gDNA extractions.

### Assay Optimization

Optimization of LAMP conditions was conducted according to the protocol of WarmStart® Colorimetric LAMP 2X Master Mix (#M1800, New England Biolabs, MA, USA). All primers were synthesized by Integrated DNA Technologies (IA, USA). The basic reaction system contains 2× WarmStart Colorimetric LAMP 2X Master Mix, primer set (the ratio of outer primers, inner primers, and loop primers was 1:4:2), gDNA or dH_2_O template, and dH_2_O. The basic incubation time was set to 30 min at 65°C. For the optimization, we used the selected primer set to run the positive samples (*B. mandrillaris* gDNA) and negative controls (dH_2_O water) under different conditions: (i) various primer concentrations at 0.75×, 1×, 1.25×, 1.5× and 1.75×; (ii) different temperatures at 57°C, 59°C, 61°C, 63°C, 65°C, and 67°C. Amplification outcomes were assessed by visual inspection of colorimetric change and by 1.5% agarose gel electrophoresis. Optimal conditions were defined as those yielding a clear pink-to-yellow (negative-to-positive) colorimetric change and characteristic ladder-like LAMP amplification patterns in positive samples, with no amplification observed in the negative control. After selecting the optimal primer concentration and temperature, we compared the amplification efficacy at different times (10-, 20-, 30-, 40-, and 50-minutes) to determine the minimum and optimal reaction times.

### Determination of LAMP Specificity and Sensitivity

Using the optimized LAMP conditions established above, assay specificity was evaluated by testing gDNA from ten different *B. mandrillaris* isolates (V039, SAM, RP5, OK1, ITSON, V433, V188, V194, V426, and V619), as well as gDNA templates from other pFLAs (*A. castellanii* and *N. fowleri*), protozoa (*S. neurona* and *T. gondii*), fungi (*C. auris* and *C. albicans*), and bacteria (*E. coli, S. marcescens*, and *P. aeruginosa*). Amplification products were evaluated by both visual inspection of colorimetric changes and by electrophoresis on 1.5% agarose gels.

To evaluate the LAMP sensitivity, the limit of detection (LOD) was compared with a previously published conventional PCR method (Booton 2003). A 10-fold serial dilution series of *B. mandrillaris* gDNA (ranging from 0.1 fg/μL to 100 pg/μL) from heat-treated cells for 1 h and standard original cells (from 1,000 to 0.001 trophozoites/100 μL or 10 to 0.0001 cells/reaction) were prepared. For each template type and cell concentration, gDNA was extracted and amplification was performed in parallel using our LAMP assay and the conventional PCR assay. The lowest template concentration consistently yielding a positive result was defined as the LOD (n=3). LAMP products were evaluated by visual inspection of colorimetric changes and further confirmed by 1.5% agarose gel electrophoresis. Conventional PCR products were analyzed by 1.5% agarose gel electrophoresis only.

For conventional PCR, primers Bal18SF3-Fwd (5′-TGG TGG AGT GAT TTG TCT GG-3′) and Bal18SB3-Rev (5′-TCG GCT AAT CAC CGA TTG TC-3′) were used. The PCR cycling conditions were as follows: initial denaturation at 94°C for 5 min; 40 cycles of 94°C for 1 min, 54°C for 2 min, and 72°C for 3 min; followed by a final extension at 72°C for 10 min (Booton 2003).

### Target gene cloning and transformation

The target gene was amplified using the outer primers of the selected LAMP primer set with *B. mandrillaris* gDNA as the template. PCR products (45 μL) were mixed with 5 μL of 10× DNA loading buffer and analyzed by electrophoresis on a 1% agarose gel. PCR products yielding a distinct and intense target band were excised and purified using an agarose gel DNA extraction kit (#28706, QIAGEN, Germany) according to the manufacturer’s instructions.

Purified PCR products were ligated into the pGEM®-T Easy Vector (Promega, MI, USA) using T4 DNA ligase and subsequently transformed into J109 competent cells (*E. coli*). Transformants were plated onto LB agar plates (5 g yeast extract, 10 g peptone, 10 g NaCl, 12 g agar, with 1 L water) supplemented with ampicillin (Thermo scientific, IL, USA), IPTG (Thermo scientific, IL, USA), and X-Gal (Thermo scientific, IL, USA), and recombinant clones were identified by blue-white screening. Positive colonies were selected and expanded by SOC (super optimal broth with catabolite repression) culture (20 g tryptone, 5 g yeast extract, 0.5 g sodium chloride, 0.186 g potassium chloride, 1 M glucose, 2 M magnesium chloride, with 1 L water). The plasmid DNA was extracted by the plasmid purification kit (QIAGEN, MD, USA). The identity of the inserted target gene was confirmed by DNA sequencing (Eurofins Scientific, KY, USA). Plasmid DNA concentration was measured using a NanoDrop spectrophotometer (Thermo Scientific, IL, USA).

### qPCR and dPCR validation assay development

The qPCR assay was developed using the outer primers from the selected LAMP primer set as the qPCR primers. Recombinant plasmids containing the target sequence were used as standard templates. qPCR amplification was carried out using a SYBR Green master mix (Thermo Scientific, IL, USA), with the reaction mixture consisting of 10 μL Master Mix, 1 μL F3 (500 nM), 1 μL B3 (500 nM), 1 μL template and 7 μL dH_2_O. A two-step amplification protocol was developed and used as follows: initial denaturation at 95 °C for 5 min, followed by 35 cycles of denaturation at 95 °C for 15 s and annealing at 61 °C for 1 min. Melt curve analysis was performed with the following program: 95 °C for 15 s, 60 °C for 1 min, and 95 °C for 1 s. The developed qPCR demonstrated a high amplification efficiency and specific single-peak melting curves. A standard curve was generated by plotting the cycle threshold (Ct) values against the logarithm of the plasmid copy number. The standard curve equation was generated by linear regression analysis, and the correlation coefficient (R^2^) was calculated by Prism Ver.10 (GraphPad, US).

Recombinant plasmid standards were prepared by 10-fold serial dilutions, and six concentrations were selected as templates for dPCR analysis to determine the absolute copy number. Each concentration was tested in triplicate, with nuclease-free dH_2_O water included as a no-template negative control. Each dPCR reaction was performed in a total volume of 40 μL in QIAcuity Nanoplate 26k (24-well) (QIAGEN, MD, USA), containing 13.3 μL of 3× EvaGreen PCR Master Mix (QIAGEN, MD, USA), 0.8 μL of F3 and B3 primers (20 μM), 1 μL of serially diluted plasmid template, and 24.1 μL of nuclease-free dH_2_O water. The amplification program was same as the qPCR protocol described above, consisting of an initial denaturation at 95 °C for 30 s, followed by 35 cycles of denaturation at 95 °C for 15 s and annealing at 61 °C for 1 min. Plates were read in a QIAcuity One dPCR system (QIAGEN, MD, USA) using the green fluorescence channel (with an excitation and emission peak of EvaGreen at 495 nm and 530 nm) with an exposure duration of 200 ms and a gain of 6. A standard curve was generated by plotting the cycle threshold (Ct) values against the logarithm of the plasmid copy number from dPCR. The standard curve equation was generated by linear regression analysis, and the correlation coefficient (R^2^) was calculated by GraphPad Prism Ver.10 (MA, USA).

## Results

### Primer Design

LAMP primers were designed to target and amplify the *B. mandrillaris* 18S rRNA gene (GenBank accession no. AF477019-AF477022, AF019071, JX524850). Multiple sequence alignments were performed to identify conserved regions across the selected sequences (http://multalin.toulouse.inra.fr/multalin/), and candidate LAMP primer sets were designed using PrimerExplorer V5 (https://primerexplorer.jp/lampv5e/index.html). The LAMP primer sets contain two inner primers (FIP and BIP) and two outer primers (F3 and B3). After the preliminary primer screening, two loop primers (LF and LB) were designed based on the selected inner and outer primers. All the candidate primers were first evaluated for sequence specificity by BLAST analysis against the NCBI database (https://www.ncbi.nlm.nih.gov/). The primers were then analyzed using OligoAnalyzer™ (Integrated DNA Technologies, USA) to evaluate stability, secondary structure formation, and dimerization. The sequences and positions of the primers are shown in **Table 1** and **Figure 1**.

**Table 1.**
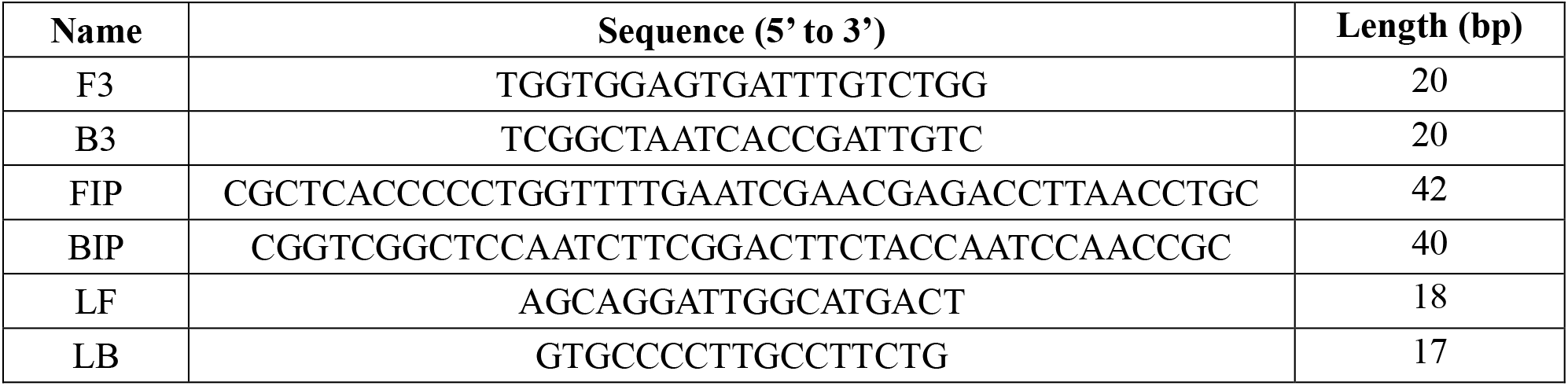
Primer sequences for LAMP diagnostic assay.

**Figure 1.**
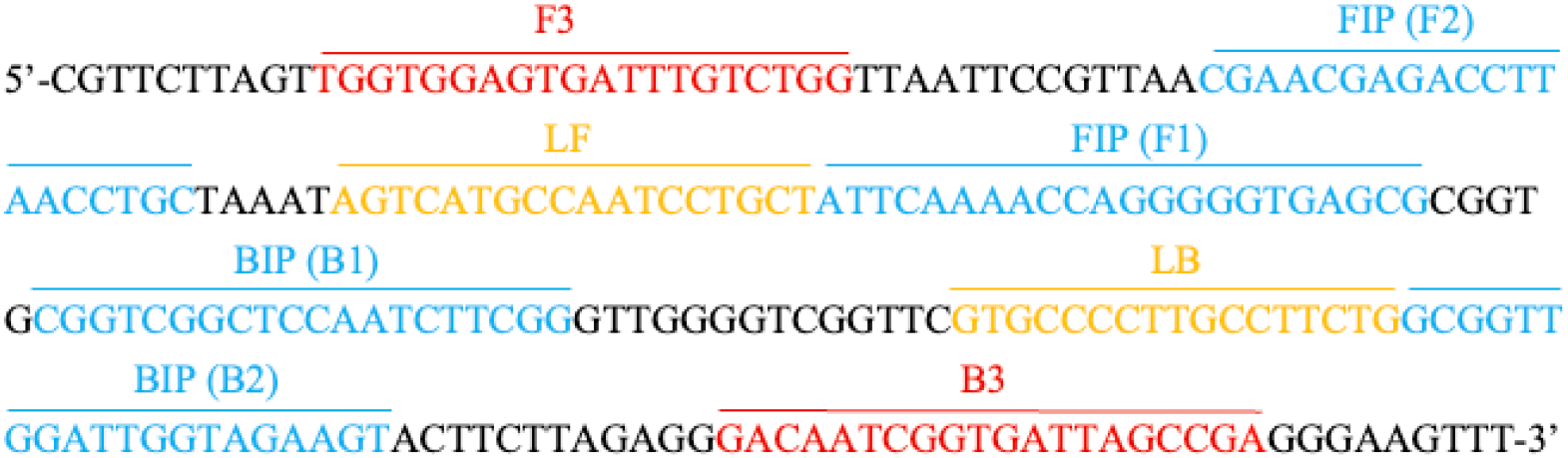
The positions of LAMP primers targeting *B. mandrillaris* 18S rRNA.

### Optimization of the LAMP assay

To optimize the LAMP amplification conditions, primer concentrations ranging from 0.75× to 1.75× were evaluated across a temperature gradient of 57-67°C. Amplification efficiency was assessed by both colorimetric readout and agarose gel electrophoresis (**Figure 2**). Overall, amplification efficiency increased with increasing reaction temperature. At 57°C, although the amplification could be observed on agarose gels, positive samples remained orange rather than a clear yellow color, that was difficult to distinguish from negative controls. Primer concentration also markedly influenced assay performance. At higher temperatures, some false-positive amplifications were observed at high primer concentrations (1.5× and 1.75×). In contrast, reactions containing 0.75× primer showed insufficient amplification with an unclear color change. Based on these observations, a primer concentration of 1× (1.28 µM FIP and BIP, 0.16 µM F3 and B3, 0.32 µM LF and LB) provided optimal assay performance. Under these constrained conditions, positive samples consistently exhibited a clear color change to yellow, while negative samples remained pink across temperatures from 59 to 67 °C. Importantly, no false-positive ladder-like bands were observed in negative controls. Based on these results, 65 °C was selected as the optimal reaction temperature, as it provided robust amplification of positive samples with high specificity.

**Figure 2.**
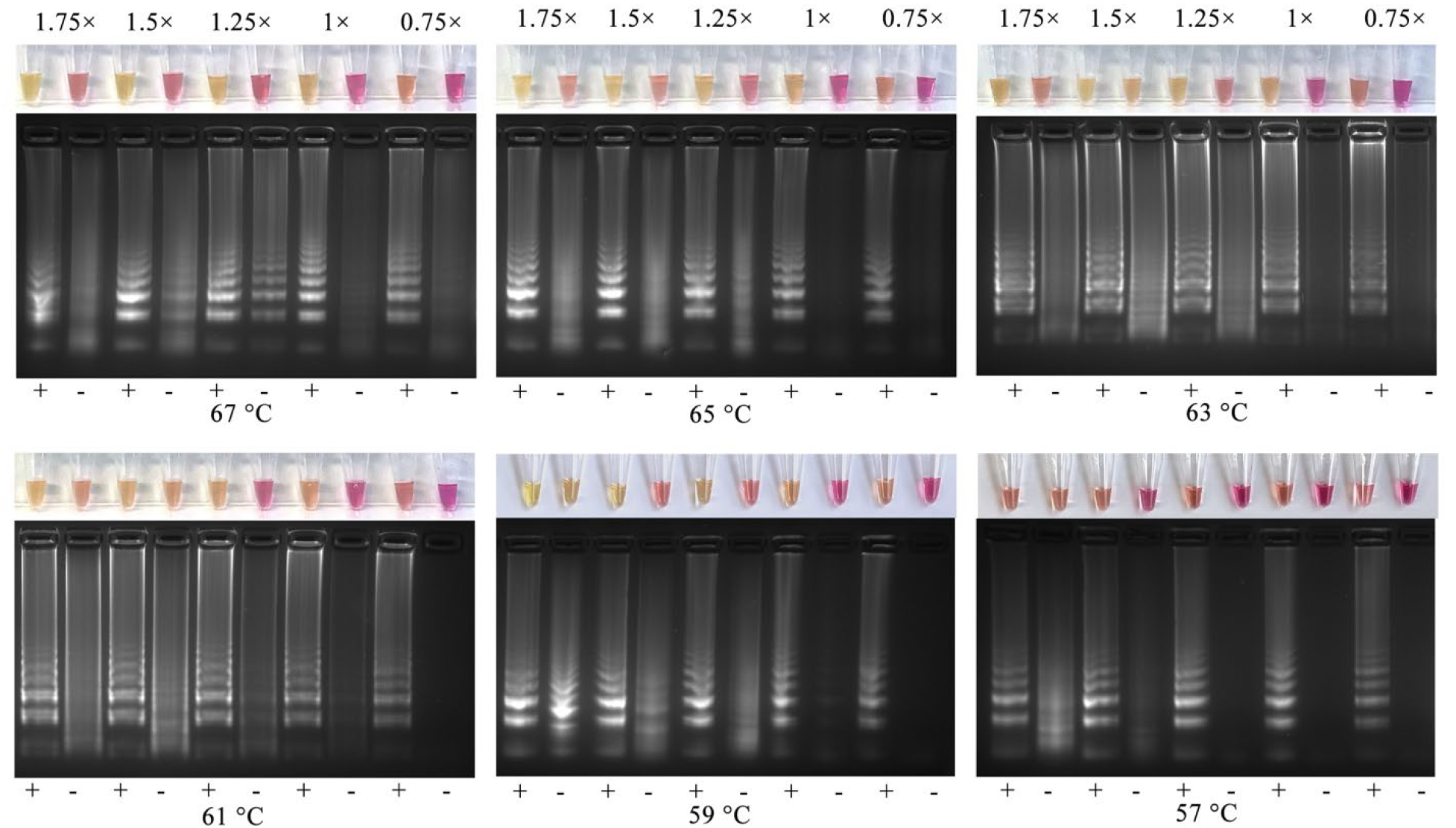
Optimization of primer concentrations at specific temperatures (57-67 °C).

After determining the optimal reaction temperature and primer concentration, the amplification time was further optimized using the selected conditions. LAMP reactions were terminated at different time points. As shown in **Figure 3**, amplification was first detectable at 20 min by the weak colorimetric orange colour, instead of yellow, and a ladder-like bands on the gel electrophoresis. At 30 min, robust amplification was observed. In contrast, reactions extended to 40- and 50-min resulted in nonspecific amplification; at 40 mins there was a slight colorimetric change and a band/streak from the negative control reaction. In the 50 min reaction, a change in colour to orange was observed and a false-positive ladder-like bands were observed in negative controls. Based on these results, a strict reaction time of 30 min was selected as the optimal amplification duration for our 18S rRNA template LAMP assay.

**Figure 3.**
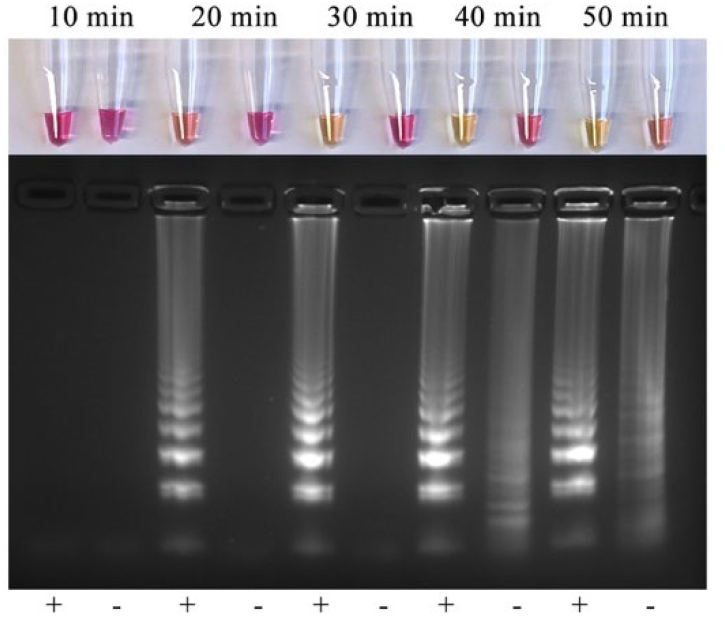
Optimization of amplification time from 10- to 50-minutes.

### The Specificity of LAMP Assay

Amplification occurred exclusively in *B. mandrillaris* isolates (reaction 1-10), with a distinct color change and the presence of ladder-like bands. In contrast, no colorimetric change or ladder-like bands were observed in non-*B. mandrillaris* organisms or molecular grade dH2O water (**Figure 4**). These results demonstrate that our novel 18S rRNA LAMP assay exhibits high specificity for *B. mandrillaris*, with no detectable cross-reactivity against other FLA, protozoa, fungal or bacterial species tested; which we had access to.

**Figure 4.**
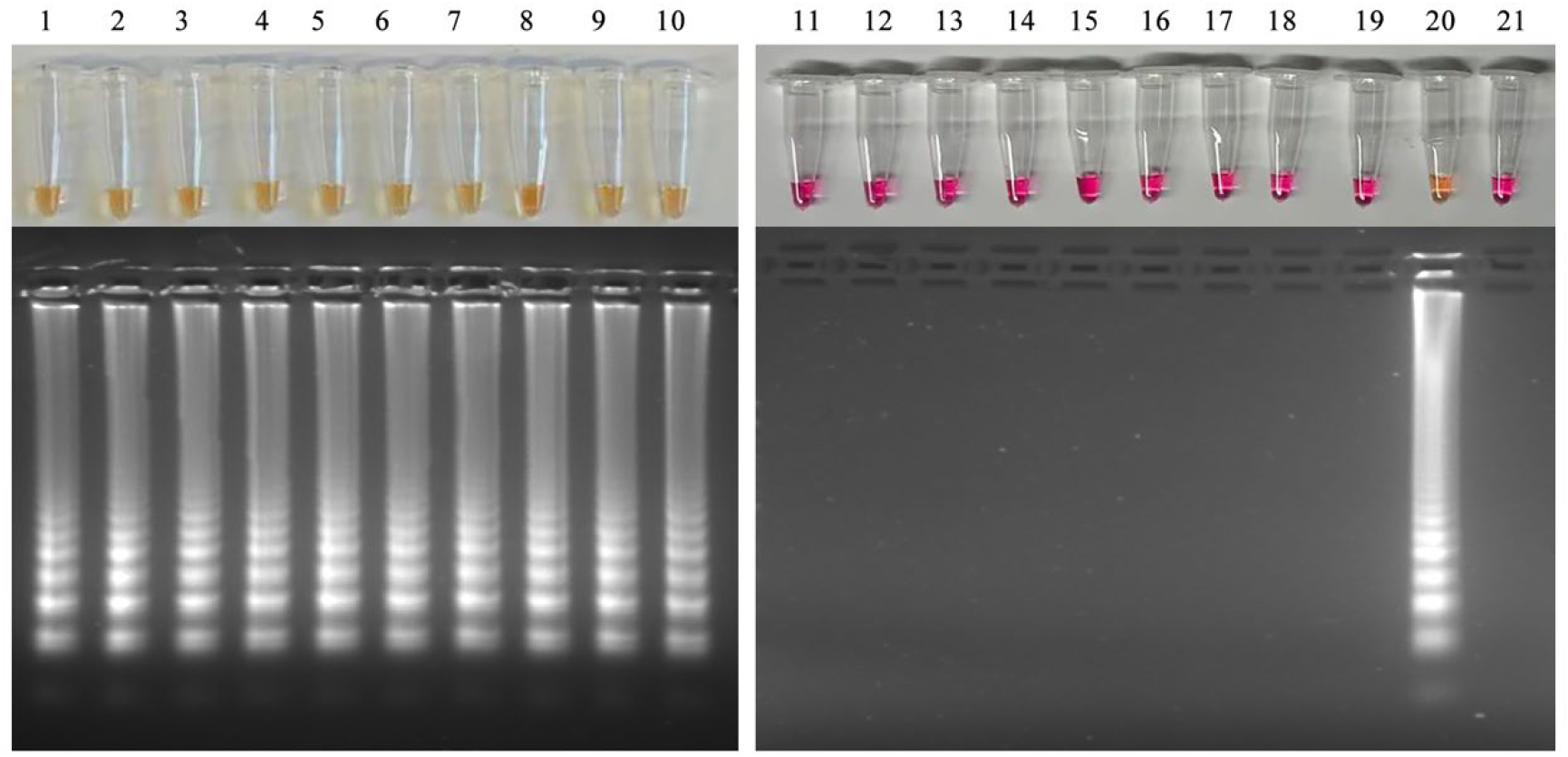
Specificity of the 18S rRNA LAMP assay. Colorimetric LAMP amplification and agarose gel electrophoresis were performed to evaluate assay specificity. Lanes 1-10 represent different *B. mandrillaris* isolates: V039, SAM, RP5, OK1, ITSON, V433, V188, V194, V426 and V619; Lane 11: *Acanthamoeba castellanii*; lane 12: *Naegleria fowleri*; lane 13: *Sarcocystis neurona;* lane 14: *Toxoplasma gondii*; lane 15: *Escherichia coli*; lane 16: *Serratia marcescens*; lane 17: *Pseudomonas aeruginosa;* 18: *Candida auris*; lane 19: *Candida albicans*; lane 20: positive control (*B. mandrillaris* V039, 1 ng); lane 21: negative control – molecular grade dH2O water.

### The Sensitivity of LAMP Assay

The sensitivity of the optimized LAMP assay was evaluated and directly compared with conventional PCR using different *B. mandrillaris* gDNA templates (**Figure 5**). For DNA-based sensitivity analysis, the LAMP assay successfully detected *B. mandrillaris* DNA from 100 pg/μL to 10 fg/μL. Conventional PCR produced detectable amplification products within a narrower range, from 100 pg/μL to 100 fg/μL. To further assess sensitivity using whole-cell templates, the LAMP assay detected *B. mandrillaris* at a minimum concentration of 0.1 trophozoites/μL, whereas conventional PCR required at least 1 trophozoite/μL for detection. When heat-treated cells were used as templates, the LAMP assay demonstrated improved sensitivity compared with original cells, with a detection limit of 0.01 trophozoites/μL (10-fold increase), while conventional PCR detected *B. mandrillaris* at the concentration of 1 trophozoites/μL (100-fold increase). Overall, across DNA-based, untreated and heat-treated cell-based gDNA templates, the LAMP assay consistently exhibited higher sensitivity compared with conventional PCR, with an estimated 10- to 100-fold increase in detection efficiency depending on the sample preparation method.

**Figure 5.**
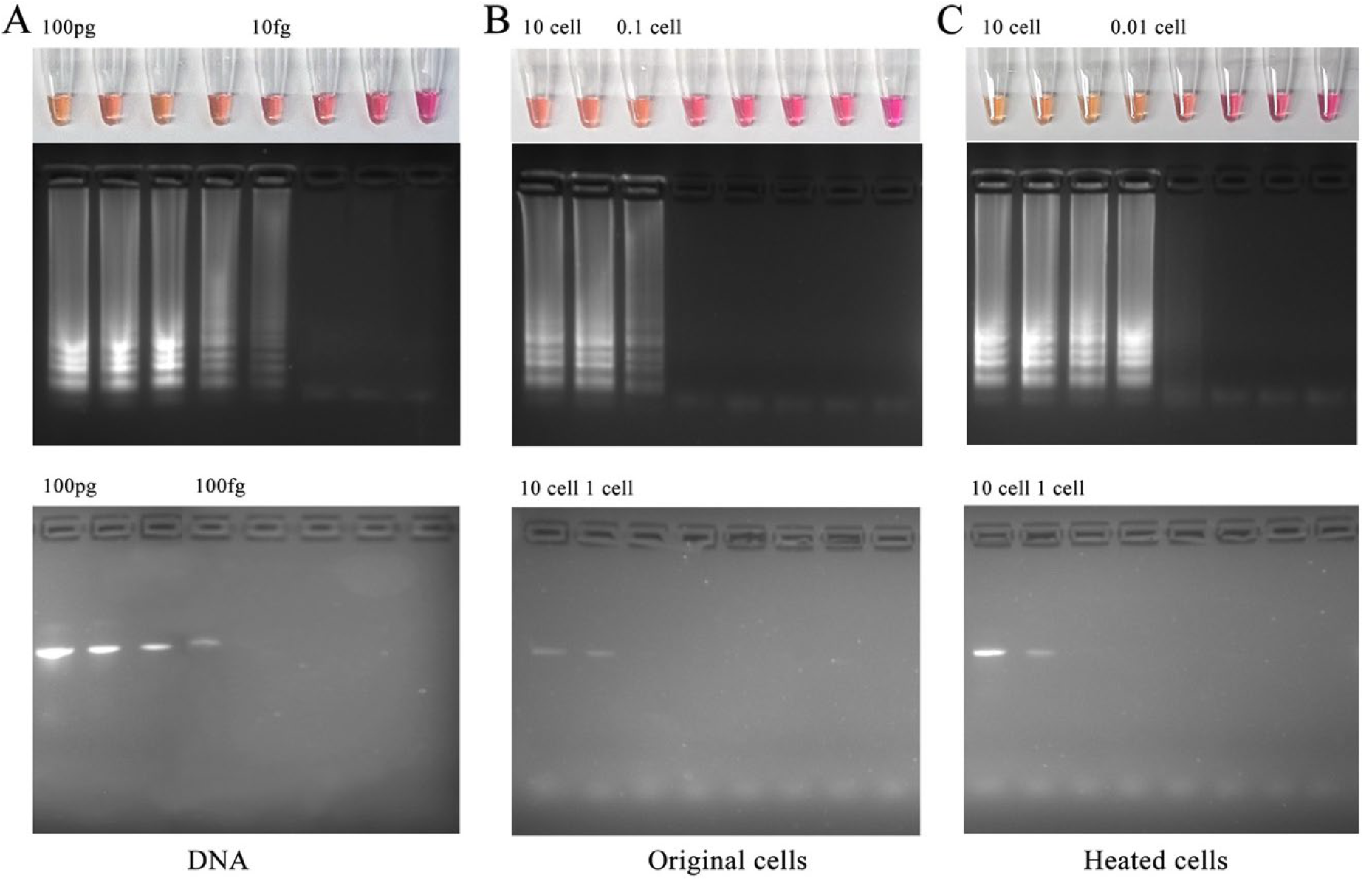
Comparison of Sensitivity of LAMP and Conventional PCR. (A) Purified *B. mandrillaris* DNA. (B) Original, untreated *B. mandrillaris* cells. (C) Heat-treated *B. mandrillaris* cells. All conditions were performed and then gDNA was extracted from each of the tested conditions and used as template for these experiments.

### Validation of LAMP Diagnostic Method

#### Copy Number Determination

To validate the LAMP assay and determine the true known limits of detection we opted to use both dPCR and qPCR protocols for quantification. The dPCR results showed a clear concentration-dependent decrease in copy number across the 10-fold serial dilution series (**Figure 6**). Clear separation between positive and negative partitions was observed in each sample, and as expected, there was no amplification from the negative control templates.

**Figure 6.**
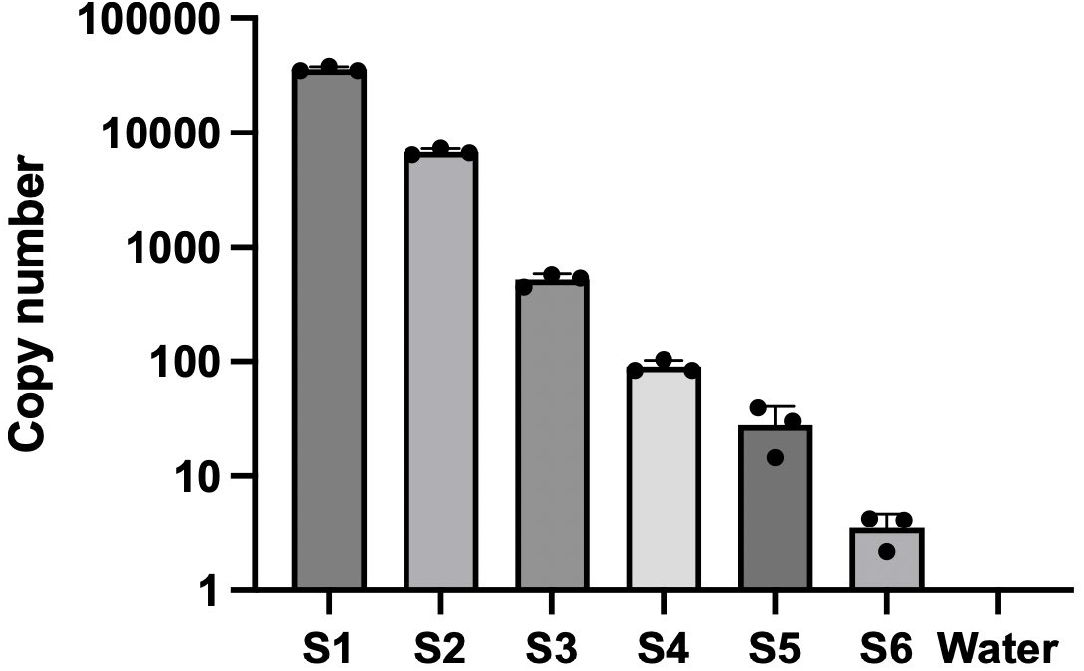
dPCR quantification of recombinant 18S rRNA plasmid standards and dH_2_O control.

Serial-diluted recombinant 18S rRNA plasmid standards with known copy numbers from the dPCR were also used as templates for qPCR amplification. The results demonstrated a strong linear relationship between the Ct values and the logarithm of plasmid copy number (**Figure 7**); while no amplification was observed in the negative control. The standard curve equation was established as Y = −3.233 x +33.91, with an R^2^ of 0.9532.

**Figure 7.**
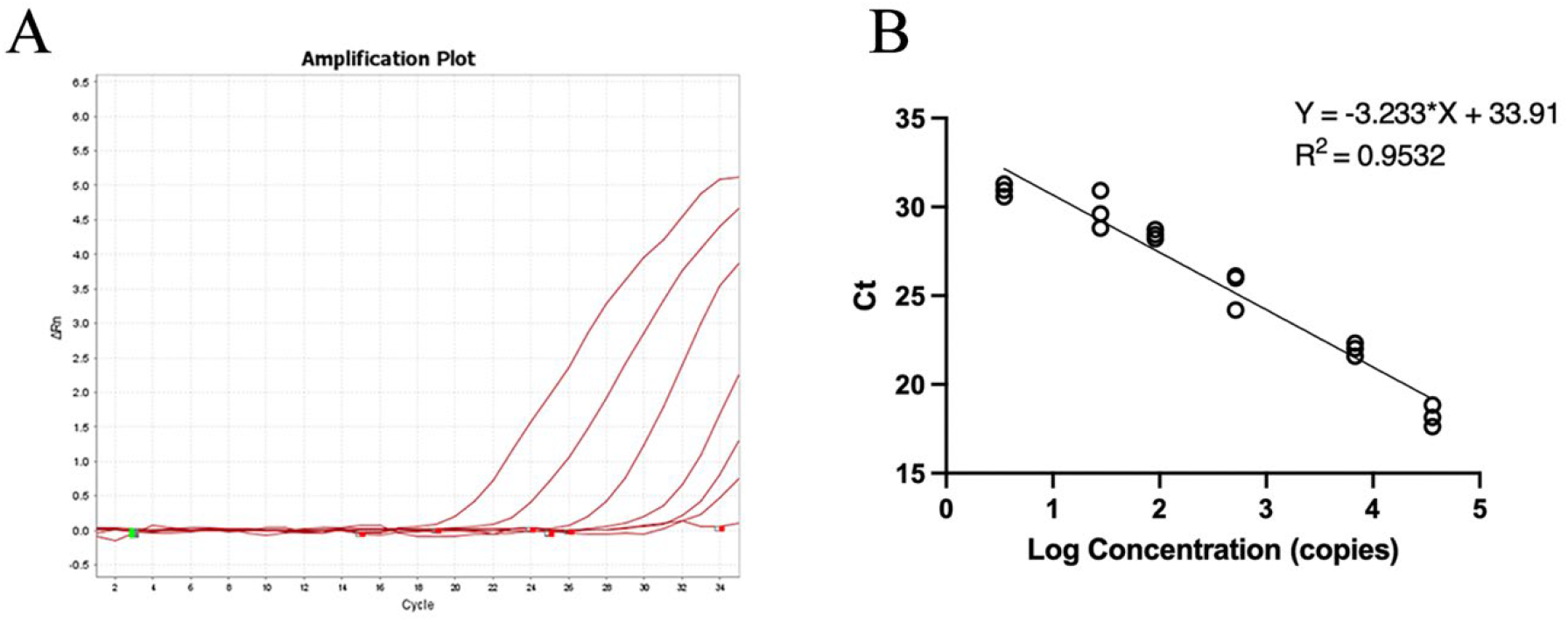
Standard Curve of qPCR A: Amplification curve of qPCR for different concentrations of *B. mandrillaris* plasmid standards and dH_2_O; B: Standard curve of qPCR results.

Furthermore, gDNA was used to evaluate the consistency between qPCR and dPCR quantification and to assess the LOD of the LAMP assay. *B. mandrillaris* gDNA samples at 1 ng/µL were quantified by both assays. The copy numbers were calculated as 1.07 × 10^5^ copies/µL and 1.13 × 10^5^ copies/µL in qPCR and dPCR, respectively, showing the consistency and reliability of these assays. Using the lowest detectable amplification DNA concentration of LAMP and the corresponding copy number derived from the standard curve, the analytical LOD of the LAMP assay was determined to be approximately 1 copy/µL.

#### Validation of Diagnostic Potential in Human Spiked Biofluid Samples

As biofluids can significantly interfere with diagnostics, we wanted to put our newly developed diagnostic tool to the test. We obtained de-identified clinical specimens of human CSF, urine, and blood from Dr. Ryan Relich previously at Indiana University School of Medicine. We performed the same standard serial-dilution of 1 to 10^6^ *B. mandrillaris* trophozoites per reaction with the optimized conditions previously standardized. The LAMP diagnostic tool successfully detected as little as 1 trophozoite per reaction in all tested human biofluid samples (yellow in all conditions, except negative template control), while PCR showed a similar result that needed a minimum of 10 trophozoites per reaction in spiked urine samples for the detection over a three-hour period vs. 30-minutes for our LAMP diagnostic tool (**Figure 8**). These results indicate that common human biological matrices, including CSF, urine, and blood, do not inhibit our LAMP amplification and diagnostic tool. The LAMP assay demonstrated robust and better performance compared to conventional PCR while providing a rapid and simple visually interpretable readout, highlighting its advantage for *B. mandrillaris* detection in clinically relevant specimens or environmental samples.

**Figure 8.**
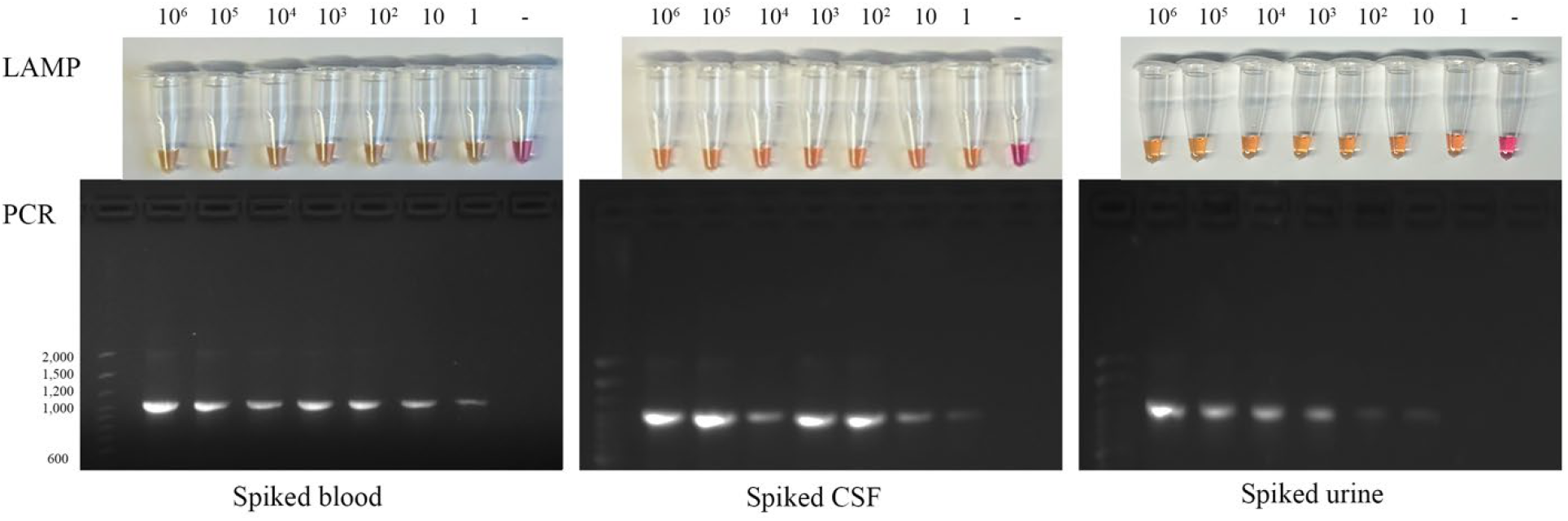
Validation of the LAMP diagnostic tool using human blood, CSF, and urine samples spiked with serially diluted *B. mandrillaris* cells, with the comparison of conventional PCR assay. Ten-fold serial dilutions, from 10^6^ to 1 trophozoite, were used as templates, and dH_2_O water was used as the negative template control.

## Discussion

The infection of *B. mandrillaris* develops rapidly once it is involved in the CNS, with the short time (weeks to months) from symptom onset to death. Early clinical manifestations are frequently subtle and nonspecific, which, together with the rarity of the disease, leads to the delay in diagnosis and timely therapeutic administration. A variety of molecular diagnostic approaches have been applied to the detection of *B. mandrillaris*. Conventional PCR assays targeting the mitochondrial 16S rRNA gene (Yagi 2005), nested PCR assays based on the 18S rRNA gene (Ahmed 2011), and real-time PCR assays targeting the ribonuclease P (RNase P) gene (Kiderlen 2008) or 18S rRNA gene (Qvarnstrom 2006) have all demonstrated variable-to-high specificity. More recently, next-generation sequencing (NGS) has emerged as a powerful tool for unbiased pathogen detection and has successfully identified *B. mandrillaris* from brain biopsy specimens previously (Wu 2020). However, these diagnoses remain limited by high cost, long times, specialized and trained staff, and the possibility of false-negative results. Compared with conventional PCR, LAMP has several advantages, like higher sensitivity, shorter reaction time, and minimal equipment requirements, making LAMP particularly suitable for use in resource-limited areas and could provide an initial primary diagnosis to start treatment sooner. Therefore, we developed the first and new diagnostic method, loop-mediated isothermal amplification (LAMP), that is cheap, quick, specific, and highly sensitive for *B. mandrillaris* diseases.

Primer design is a critical determinant of LAMP assay performance. Because we obtained false-positives from various other species when we tried to target *B. mandrillaris* 16S rRNA, and that the18S rRNA gene had been previously reported used as targeted gene in previous PCR-based diagnostics (Qvarnstrom 2006, Ahmed 2011); this study selected and focused on the 18S rRNA gene. In addition to primer design, other parameters, including temperature, reaction time, dye used for colorimetric change, and primer concentration were also optimized, as these factors significantly affect amplification efficiency and visual readout/change. The LAMP assay contains six primers, so the reaction system is easy to cause primer-primer interactions, forming secondary structures, which can lead to nonspecific amplification and false-positive results. In this study, both positive and negative controls were included for each tested condition during the optimization, which allowed reliable discrimination between true target-specific amplification and nonspecific background signals, thereby minimizing the risk of false-positive interpretation and ensuring robust assay performance. Under optimized conditions, the established LAMP assay showed a rapid amplification at 65 °C within 30 min, with primer concentration of 1.28 µM FIP and BIP, 0.16 µM F3 and B3, 0.32 µM LF and LB, exhibiting high specificity without cross-reactivity to other amoebas (highlighting no cross reactivity with *Acanthamoeba* as this has been seen before), protozoa, fungi, or bacteria used within this study. Next we would like to expand our testing to other CNS/meningitis causing pathogens.

Another important consideration in LAMP assays is the sample preparation. The traditional molecular diagnostic assays typically require nucleic acid extraction and purification, which increases cost and time. One of the major strengths of LAMP is its tolerance to crude samples (Soroka 2021). Accordingly, this study compared multiple sample preparation methods, including commercial DNA extraction kits, rough extraction (heating), and direct use of samples. The results demonstrated that the developed LAMP assay can detect *B. mandrillaris* from the rough extraction samples, even the original samples, highlighting its robustness and suitability for rapid testing workflows. Furthermore, the comparison with conventional PCR demonstrated that our LAMP diagnostic tool exhibits a 10- to 100-fold higher amplification efficiency, which enables reliable detection of very low numbers of *B. mandrillaris* clinically. This study is the first to develop, validate and describe a dPCR assay against any pFLA. We used this to determine the copy number of the *B. mandrillaris* 18S rRNA target with LOD and further validated this using our novel developed qPCR method also. The LAMP diagnostic tool was then evaluated by copy number, which allowed accurate calculation of the limit of detection. This quantification provides a significant advantage over qualitative detection alone, as it can provide direct proof to support standardized assessment of assay sensitivity and performance.

Although there are many positives of using LAMP based diagnostics, some of the challenges in LAMP assays is contamination due to the large quantity of amplification products generated in a short time and the potential formation of aerosols. In our study, phenol red was added before amplification, allowing results to be interpreted directly by the naked eye without opening reaction tubes. However, in practical applications, colorimetric interpretation may introduce observer-dependent variability, especially under different lighting conditions or when weak positive reactions produce borderline color changes (orange colour change). Agarose gel electrophoresis may still be required to confirm ambiguous results. This additional step thereby increases the risk of amplicon contamination and potentially leads to false-positive results in subsequent assays. Therefore, LAMP results should be considered as a rapid preliminary diagnostic readout, and other diagnostic methods can be used for further confirmation and validation of *B. mandrillaris* infection.

## Conclusion

This study developed and validated three novel diagnostic methods (LAMP, qPCR, and dPCR) for *B. mandrillaris*. By optimizing primers, primer concentrations, and reaction conditions, specificity and sensitivity determination, copy number quantification, and spiked human biofluid sample validation, we demonstrated that the LAMP assay has high sensitivity, specificity, and robustness, which make this method a future diagnostic tool well suited for early clinical diagnosis, especially in primary care facilities or resource-limited areas. Early diagnosis may facilitate timely treatment, ultimately improve patient outcomes, and in particular decrease the high mortality rate associated with BAE and other *Balamuthia* diseases.

## Supplemental material

A primer set targeting the 16S rRNA gene was also designed and evaluated during assay development; the sequences were shown in **Table S1**. Specificity testing revealed cross-reactivity with *S. neurona* and bacteria (*E. coli, S. marcescens*, and *P. aeruginosa*), as shown in **Figure S1**. Consequently, this primer set was excluded from further evaluation, and the 18S rRNA gene was selected as the target for the final assay.

**Figure S1.**
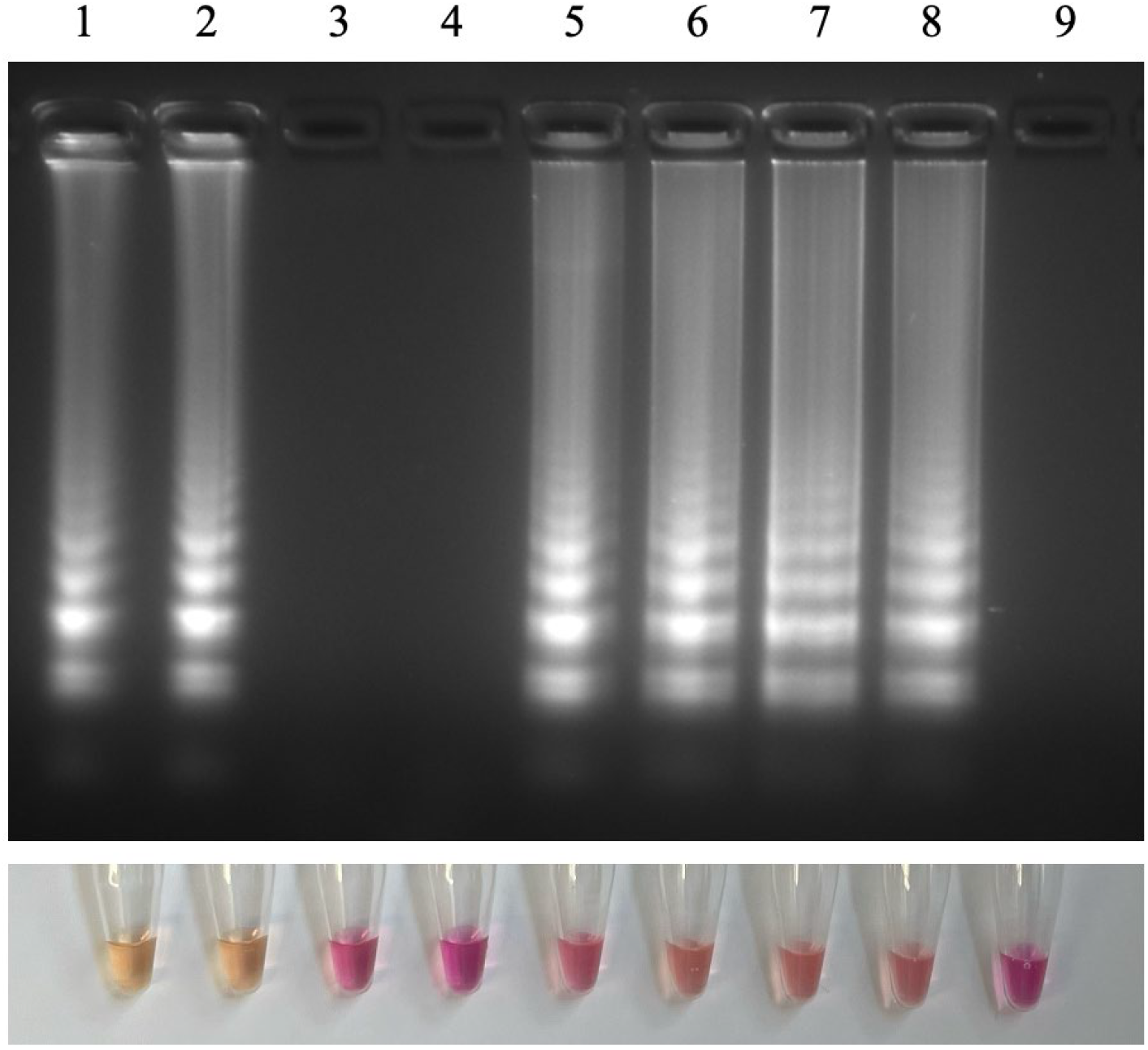
Specificity of LAMP targeting 16S rRNA. Lane 1: *B. mandrillaris* (V039); lane 2: *B. mandrillaris* (ITSON); lane 3: *N. fowleri*; lane 4: *A. castellanii*; lane 5: *S. neurona*; lane 6: *E. coli*; lane 7: *S. marcescens*; lane 8. *P. aeruginoscreening. Positive colonies were selectedsa*; lane 9: negative control – molecular grade dH_2_O water.

**Table S1.**
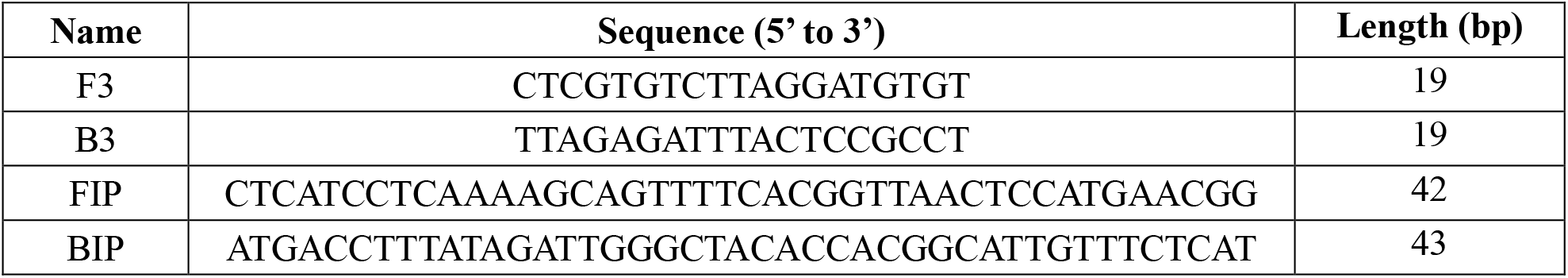
Primer sequences targeting 16S rRNA.

